# Reciprocal targeting of the unfolded protein response regulator Xbp1 and the Dom-A nucleosome remodeler in *Drosophila*

**DOI:** 10.1101/2025.10.15.682518

**Authors:** Gizem Kars, Peter B. Becker, Zivkos Apostolou

## Abstract

The DOM-A complex regulates cell growth and proliferation in *Drosophila*. Like the orthologous human P400 complex, DOM-A combines two epigenetic effectors: a SWR1-type histone exchange enzyme, Dom-A, and the Tip60 acetyltransferase. We found Xbp1, a conserved transcription regulator of the unfolded protein response (UPR), as tightly associated with immunopurified DOM-A and explored the functional implications of this interaction.

We biochemically determined the Xbp1 DNA recognition motif in chromatin-reconstituted *Drosophila* genomes. Intersection of the chromatin binding profiles for Xbp1 and Dom-A in proliferating cells and reciprocal protein depletion studies revealed two distinct modes through which Xbp1 binds chromatin. Xbp1 recruits Dom-A to motif-bearing promoters of genes involved in the UPR, such as *Xbp1, Hsc70-3* and *Gp93*, and activates their transcription. Xbp1 also localizes to hundreds of high-confidence Dom-A binding sites that lack Xbp1 recognition motifs. These interactions depend on Dom-A, pointing to a ‘reverse targeting’ scenario. Upon depletion of Dom-A, Xbp1 protein levels, but not mRNA levels, are reduced. The Xbp1 may thus be stabilized upon binding to DOM-A. The complex interactions of Xbp1 and DOM-A in the genome bear potential to integrate signals from the UPR with the general, DOM-mediated regulation of cell growth and proliferation.

**Graphical abstract:** 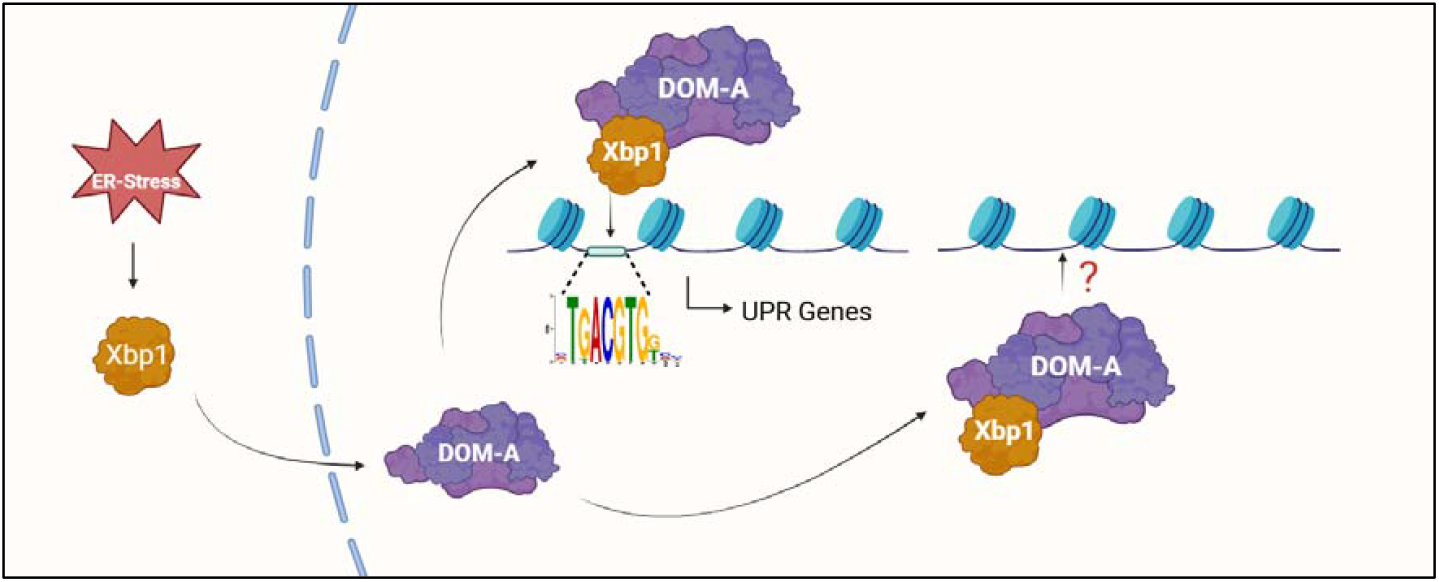

## Introduction

Epigenetic regulators of gene expression are often targeted to their chromosomal sites of action by sequence-specific DNA-binding transcription factors (TF) to modulate local chromatin organization (1). One potent regulator is the large, 15-subunit Domino-A (DOM-A) complex of *Drosophila*, which combines acetyltransferase and nucleosome remodeling activities (2). Dom-A, the signature subunit of the DOM-A complex, is related to the SWR1 nucleosome remodeling enzyme in yeast (3, 4). ATPases of this type exchange the regular histone H2A with variants to produce nucleosomes with special properties (5, 6). The Tip60 acetyltransferase subunit acetylates the histone H4 and the variant H2A.V (H2A.Z in mammals) at the first nucleosome downstream of transcription start sites, and this acetylation correlates with transcriptional activity (7).

The DOM-A complex is required for efficient cell growth and proliferation (8) It localizes to many active gene promoters in proliferating *Drosophila* cells, notably of genes coding for proteins involved in various aspects of the cell cycle, from replication to mitosis (7). The subunits of the Dom-A complex have been conserved during evolution. The orthologous regulator in mammals is termed the p400/TIP60 complex (2). Like DOM-A, p400/TIP60 complexes are involved in regulating cell proliferation (9–15).

How are DOM-A complexes recruited to their sites of action? Some subunits of DOM-type complexes bear bromo and YEATS domains that may interact with acetylated nucleosomes around transcription start sites(4, 16, 17). SWR1-type ATPases are also attracted to nucleosome-free regions (NFRs) that characterize active promoters (18). Histone acetylation and NFRs are generic features of active promoters, and more selective targeting involves interaction with sequence-specific TFs. Studies in mammals have shown that p400/TIP60 complexes are recruited by TFs to activate specific gene promoters, frequently via interactions with the large TRRAP subunit (19–22). Interaction between TFs and Domino Complexes has also been identified before (23–25)

For the *Drosophila* Dom-A complex, no targeting cues have been identified so far. However, when we characterized the DOM-A complex by immunopurification and mass spectrometry (IP-MS), we found the transcription factor Xbp1 (X-box binding protein 1) tightly associated, along with *bona fide* subunits (3). This finding was particularly striking, as the purification was performed under stringent conditions using a combination of detergents and benzonase to eliminate DNA-mediated interactions, thereby minimizing non-specific protein associations. These conditions support the conclusion that Xbp1 is tightly associated with the core DOM-A complex. We hypothesized that the interaction of Xbp1 would target the Dom-A complex to particular promoters.

*Drosophila* Xbp1 is one of three transcription factors (Atf4, Atf6, Xbp1) that mediate transcriptional effects of the ‘Unfolded Protein Response’ (UPR). Cells can detect the accumulation of unfolded proteins that arises from environmental stress, such as heat, or from defects in protein folding in the endoplasmic reticulum (ER). In response, the UPR is activated, implementing several proteostasis mechanisms, including reducing overall gene expression, enhancing protein folding capacity and promoting the degradation of misfolded proteins (26, 27).

Xbp1 is activated in one of three evolutionary conserved ‘arms’ of the UPR, involving the kinase Ire1 (inositol-requiring kinase). Ire1 is an ER transmembrane protein. In the presence of unfolded peptides, binding of the endoplasmic heat shock protein BiP (Hsc70-3 in *Drosophila*) to the luminal side of Ire1 triggers the dimerization and autophosphorylation of the kinase. This activates an endonuclease activity that removes a small intron from *Xbp1* mRNA, which is now translated into the active Xbp1 transcription factor (28).

In addition to its role in the UPR, Ire1 may also be activated independently of ER stress in the context of extracellular signaling pathways. Accordingly, Xbp1 regulates diverse aspects of cell differentiation, tissue homeostasis and metabolism suggesting complex intersections of the UPR with developmental mechanisms (28–33). The developmental loss-of-function phenotypes can be attributed to the protein’s essential role in tissues with high secretory demand (33–36).

Motivated by the discovery of the strong physical interaction of Xbp1 with the DOM-A coactivator, we explored the functional implications in our cell system. We defined Xbp1’s binding motif and its potential binding sites in the genome by an *in vitro* ChIP approach. We established the first genome-wide binding profile using the CUT&RUN methodology and intersected it with Dom-A as well as Tip60 profiles. We observed that Xbp1 recruits the Dom-A complex to a set of genes involved in the UPR. Surprisingly, we also observed a ‘reverse targeting’ scenario. We detected low levels of Xbp1 binding at many Dom-A binding sites, most of which lack an Xbp1 consensus sequence. Dom-A depletion results in reduced levels of Xbp1 and strongly reduced promoter binding of the transcription factor. We speculate that the interaction between Xbp1 and Dom-A stabilizes the transcription factor, which could account for the initial identification of Xbp1 as a stable component of the DOM-A complex. Conversely, Dom-A appears to be a critical requirement for Xbp1 binding at key UPR target genes, such as *Xbp1* and *Hsc70-3*.

## Materials and Methods

### Cell lines and culture conditions

*Drosophila* L2-4 cells (S2 subclone, kind gift from Patrick Heun) were cultured at 26□°C in Schneider’s medium (Thermo-Fisher, Cat. No. 21720024) supplemented with 10% FBS (PAN Biotech, Cat. No. P30-3031) and 1% Penicillin-Streptomycin (Sigma-Aldrich, Cat. No. P-4333).

### RNA interference (RNAi)

Primer sequences were obtained from DRSC/TRiP (https://fgr.hms.harvard.edu/fly-in-vivo-rnai) (Table 1). PCR products with T7 promoters were amplified from *Drosophila* genomic DNA using NEB Taq (Cat. No. M0267) in ThermoPol buffer, verified by agarose gel, and purified with the NucleoSpin Gel and PCR Clean-up Kit (Macherey-Nagel, Cat. No. 740609.50). RNA was synthesized from 1 μg PCR product using HiScribe T7 High Yield RNA Synthesis Kit (NEB, Cat. No. E2040), DNase-treated, purified by LiCl precipitation, and annealed by heating to 85°C for 5 min followed by slow cooling. One million cells per well were seeded in 6-well plates with 2 mL complete medium. The next day, medium was replaced with 500 μL serum-free Schneider’s medium, 6 μg dsRNA was added, incubated 1 h at 26°C, then 2 mL complete medium was restored. Cells were split after 3 days, and a second RNAi treatment was performed for 4 more days.

### RNA isolation, cDNA synthesis, and Real-time quantitative PCR (RT-qPCR)

Five million L2-4 cells were collected, washed with PBS and total RNA extracted using the RNeasy Kit (Qiagen, Cat. No. 74104). RNA concentration was measured by NanoDrop, and 2 μg RNA was treated with DNase I (Roche, Cat. No. 04716728001). One microgram of DNase-treated RNA was used for cDNA synthesis with SuperScript III and random hexamers (Invitrogen, Cat. No. 1080). RT-qPCR was performed with 1x SYBR Green Master Mix (Applied Biosystems, Cat. No. 4309155), 1 μM primers, and 1:50 diluted cDNA. Gene expression was calculated using the ΔΔCt method and normalized to RpII140 or 7SK. For primer sequences, see Table 1.

### Nuclear Extraction

Cells were cultured in T125 flasks, and 5×10^8^ cells were harvested (500 x g, 5 min), washed once with 10 mL PBS, and pelleted again. From this step onward, buffers were kept ice-cold and centrifugations were performed at 4°C. Cells were lysed in 10 mL NBT-10 Buffer (15 mM HEPES pH 7.5, 15 mM NaCl, 60 mM KCl, 0.5 mM EGTA, 10% sucrose, 0.15% Triton X-100, 0.2 mM PMSF, 1 mM DTT, cOmplete EDTA-free Protease inhibitor) for 10 min with rotation. Lysates were layered onto 20 mL NB-1.2 buffer (same as NBT-10 but with 1.2 M sucrose) and centrifuged (4000 rpm, 15 min) to pellet nuclei. Pellets were washed with 1 mL NB-10 Buffer (same as NBT-10 but without detergent) and centrifuged (5000 x g, 5 min). Nuclei were lysed in 500 μL SENE-500 buffer (25 mM HEPES pH 7.9, 500 mM KCl, 2 mM MgCl_2_, 0.1 mM ZnCl_2_, 0.05% IGEPAL, 0.2 mM PMSF, 1 mM DTT, cOmplete EDTA-free Protease inhibitor) on ice for 1 h, then centrifuged (16,000 x g, 15 min). The supernatant (nuclear extract) was collected, and protein concentration was determined by Bradford assay (Bio-Rad, Cat. No. 5000006) with BSA standards (Thermo Fisher, Cat. No. 23208).

### Co-immunoprecipitation (Co-IP)

All buffers were kept ice-cold, and centrifugations were performed at 4°C. For each condition, 2 mg of nuclear lysate was diluted in SENE Buffer (without KCl) to reach a final concentration of 130 mM KCl. Lysates were clarified by centrifugation at 16,000 x g for 15 min, and 40 μL of Protein A/G beads (50% slurry, equilibrated with SENE-130 Buffer) were added for pre-clearing. Samples were rotated with beads for 1 hour, followed by centrifugation to pellet the beads. The pre-cleared lysates were transferred into fresh tubes, and 40 μL of each sample was reserved as input (mixed with 10 μL of 5x Laemmli Buffer (250 mM Tris-HCl pH 6.8, 50% glycerol (v/v), 10% SDS (w/v), 0.05% Bromophenol Blue(w/v), 0.5 M DTT). For immunoprecipitation, 5 μL of anti-Xbp1 serum (Rabbit SA1228), along with corresponding pre-immune sera as controls, were added to the pre-cleared lysates. Samples were incubated overnight on a rotating wheel. The next day, 40 μL of Protein A/G beads (50% slurry, equilibrated with SENE-130 Buffer) were added to each sample, and mixtures were rotated for 3 hours. Beads were then pelleted and washed three times with SENE-150 Buffer. A final wash was performed with TBS, and beads were resuspended in 50 μL of Laemmli buffer. Unbound fractions were also collected for analysis alongside input and IP samples by western blot. For a list of antibodies, see Table 2.

### Immunoblotting

Samples in Laemmli Buffer were boiled at 95°C for 5 min and loaded onto 8% or 10% SDS-PAGE gels (Serva, Cat. Nos. 43260.01, 43264.01, 43270.01). Electrophoresis was carried out in Running Buffer (25 mM Tris, 192 mM glycine, 0.1% SDS) and proteins were transferred to nitrocellulose membranes (Cytiva, Cat. No. GE10600002) using wet transfer (350 mA, 1 h, 4°C). Transfer Buffer (25 mM Tris, 192 mM glycine) was supplemented with 10% methanol + 0.1% SDS (for Domino) or 20% methanol (for Xbp1). Membranes were blocked in 5% BSA or 5% non-fat milk in 1x TBST (5 mM Tris-HCl pH 7.5, 150 mM NaCl, 0.1% Tween-20) for 1 h at room temperature (RT), then incubated overnight at 4°C with primary antibodies in TBST containing the same blocker. After three TBST washes, membranes were incubated with LI-COR IRDye secondary antibodies (1 h, RT), washed again, and visualized on the LI-COR Odyssey CLx Imaging System. For a list of antibodies, see Table 2.

### Recombinant protein expression and purification

The spliced Xbp1 (Xbp1s) coding sequence was cloned into pETM11 with an N-terminal His-and Flag-tag and a C-terminal Spy2-tag; an untagged version was used to generate anti-Xbp1 antibodies in rabbit and guinea pig (Eurogentec). Constructs were transformed into BL21 Gold *E*.□*coli* and plated on LB-Kanamycin (50□μg/mL). Colonies were grown overnight in 10□mL LB-Kanamycin at 37°C, then 500□mL culture was inoculated (1:100) and grown at 37°C to OD_600_ ≈ 0.4. After chilling on ice for 15□min, expression was induced with 1□mM IPTG, and cultures were incubated overnight at 18°C. Cells were harvested (4000□rpm, 10□min, 4°C), washed with PBS, and lysed in 25□mL Lysis buffer (50□mM HEPES pH□7.5, 500□mM NaCl, 0.5% sarkosyl, freshly added 2□mM DTT, 0.2□mM PMSF, cOmplete EDTA-free Protease inhibitor) by sonication (6□min active; 20% output; 5□s on/off). Lysates were clarified (14,000□rpm, 15□min, 4°C) and diluted 1:1 with Binding Buffer (50□mM HEPES pH□7.5, 500□mM NaCl, 20□mM imidazole, 0.25% sarkosyl, 2□mM DTT, 0.2□mM PMSF, cOmplete EDTA-free Protease inhibitor). His-tagged Xbp1 was purified on Ni-NTA resin (Merck, Cat. No. 70666) pre-washed with Base buffer (50□mM HEPES pH□7.5, 500□mM NaCl), incubated 2□h at 4°C, washed three times with 10□mL Binding buffer, transferred to gravity columns, and eluted in eight 500□μL fractions with Elution buffer (50□mM HEPES pH□7.5, 300 mM NaCl, 250□mM imidazole). Fractions were analyzed by SDS-PAGE and Coomassie staining (Bio-Rad, Cat. No. 4568035) and protein concentration determined using a BSA standard curve (Thermo Fisher, Cat. No. 23208).

### *Drosophila* preblastoderm embryo extract (DREX)

Protocol from (37) followed for embryonic extract preparation. Oregon-R embryos were collected every 90 min until 50 mL settled volume was obtained, storing interim collections in ice-cold embryo wash (EW) buffer (0.7% NaCl, 0.04% Triton X-100). Embryos were dechorionated in 200 mL EW buffer containing 60 mL 13% sodium hypochlorite (VWR, Cat. No. APPC213322.1214) for 3 min at RT on a magnetic stirrer, rinsed under cold tap water for 5 min, and transferred to EW buffer. Floating chorions were aspirated, embryos settled in 0.7% NaCl, and resuspended in ice-cold EX10 buffer (10 mM HEPES pH 7.6, 10 mM KCl, 1.5 mM MgCl_2_, 0.5 mM EGTA, 10% glycerol, freshly added 1 mM DTT, 0.2 mM PMSF). Embryos in EX10 were transferred to a homogenizer tube (Schuett-Biotech, Cat. No. 3213402) and homogenized (1 stroke at 3000 rpm, 10 strokes at 1500 rpm, PTFE pestle). Homogenate volume was measured and supplemented with MgCl_2_ to 5 mM, mixed, and centrifuged at 27,000 x g for 10 min at 4°C. The supernatant (excluding lipid layer) was centrifuged at 245,000 x g for 2 h at 4°C. Clarified extracts were collected by syringe, avoiding lipid and pellet, aliquoted, snap-frozen in liquid N_2_, and stored at −80°C. Protein concentration was determined by Bradford assay (Bio-Rad, Cat. No. 5000006) using a BSA standard curve.

### DREX chromatin assembly and MNase digestion

For each batch, optimal extract volume for chromatin reconstitution was determined empirically. Briefly, 2 μg of *Drosophila* genomic DNA (S2 cells; Qiagen, Cat. No. 13343) was mixed with 80-100 μL DREX, 15 μL 10x McNAP ATP regeneration buffer (30 mM MgCl_2_, 10 mM DTT, 0.01 mg/mL creatine phosphate kinase, 0.3 M creatine phosphate, 0.03 M ATP), and EX-50 buffer (10 mM HEPES pH 7.6, 50 mM KCl, 1.5 mM MgCl_2_, 0.5 mM EGTA, 10% glycerol, freshly supplemented with 1 mM DTT, 0.2 mM PMSF, cOmplete EDTA-free Protease inhibitor) to 150 μL. Chromatin assembly was performed at 26°C with shaking (300 rpm, 4 h). Quality was assessed by agarose gel electrophoresis after MNase digestion. For MNase digestion, 150 μL assembled chromatin was mixed with 200 μL MNase mix (5 mM CaCl_2_, 2.3 U MNase [Sigma, Cat. No. N3755] in EX-50) and incubated at 26°C for 15, 60, or 120 s. At each time point, 110 μL was stopped with 40 μL 100 mM EDTA. DNA was extracted by phenol-chloroform-isoamyl alcohol, followed by chloroform, then precipitated with 2 μL glycogen (10 mg/mL), 150 μL 7.5 M ammonium acetate, and 880 μL ethanol at −20°C (1 h). Samples were centrifuged (16,000 x g, 15 min, 4°C), washed with 70% ethanol, air-dried for 5 min, and resuspended in 8 μL 1x TE (10 mM Tris-HCl pH 8, 1 mM EDTA). MNase fragmentation was analyzed on a 1.5% agarose gel with 2 μL 5x Orange-G loading dye.

### *In vitro* chromatin immunoprecipitation (ChIP)

Chromatin assembly was performed as described. After 4 h, 50 nM recombinant Flag-Xbp1-Spy was added and incubated for 1 h at 26°C; chromatin without recombinant protein served as negative control. Samples were mildly fixed with 0.1% formaldehyde for 10 min, quenched with 125 mM glycine for 5-10 min, and fragmented by 3 min MNase digestion to yield mononucleosomes. Reactions were adjusted to 1 mL with RIPA Buffer (25 mM HEPES-NaOH pH 7.6, 150 mM NaCl, 1% Triton X-100, 0.1% SDS, 1 mM EDTA, 0.1% sodium deoxycholate, freshly added 1 mM DTT, cOmplete EDTA-free Protease inhibitor). Protein A/G beads (20 μL slurry; Cytiva, Cat. Nos. 17-0618-05, 17-5280-05) were equilibrated in RIPA (3 x 5 min washes). For pre-clearing, samples were incubated with beads for 1 h at 4°C, then the beads were removed. Antibody (1 μL) was added and incubated overnight at 4°C with rotation. The following day, 20 μL equilibrated beads were added for 3 h at 4°C, then washed three times with RIPA. DNA was eluted by incubating beads in 100 μL 1x TE with 3.2 μL 10% SDS and 10 μL Proteinase K (10 mg/mL) at 56°C overnight. DNA was purified by phenol-chloroform-isoamyl alcohol extraction (Sigma, Cat. No. 516726), ethanol-precipitated, and resuspended in 30 μL 0.1x TE.

### CUT&RUN

CUT&RUN was performed as described (38). For each condition, 10□ cells were washed three times with Wash Buffer (WB: 20 mM HEPES-NaOH pH 7.5, 150 mM NaCl, 0.5 mM spermidine, cOmplete EDTA-free Protease inhibitor), centrifuging at 600 x g for 3 min. Cells were resuspended in 1 mL WB per antibody. Concanavalin A beads (10 μL/sample) were activated by two washes in Binding Buffer (20 mM HEPES pH 7.5, 10 mM KCl, 1 mM CaCl_2_, 1 mM MnCl_2_), mixed with cell suspension and incubated for 10 min at RT with rotation. Beads were collected on a magnet and resuspended in 150 μL Antibody Buffer (20 mM HEPES pH 7.5, 150 mM NaCl, 0.5 mM spermidine, 0.05% digitonin, 2 mM EDTA, 0.5 mM Na-butyrate) containing diluted antibodies, followed by incubation for 2 h at RT or overnight at 4°C with slow rotation. After two washes with Dig-Wash Buffer (DWB: 20 mM HEPES pH 7.5, 150 mM NaCl, 0.5 mM spermidine, 0.05% digitonin, cOmplete EDTA-free Protease inhibitor), beads were resuspended in 150 μL pAG-MNase solution (700 ng/mL in DWB) and incubated at 4°C for 1 h with slow rotation. After two further washes, beads were resuspended in 100 μL DWB, chilled, and digestion was initiated with 2 μL 100 mM CaCl_2_ for 30 min. Reactions were stopped with 100 μL 2x Stop Solution (340 mM NaCl, 20 mM EDTA, 4 mM EGTA, 0.05% digitonin, 100 μg/mL RNase A, 50 μg/mL glycogen) and incubated at 37°C for 30 min to release chromatin. Supernatants were collected on a magnet. Reverse crosslinking was done with 2 μL 10% SDS and 2.5 μL Proteinase K (20 mg/mL) at 50°C for 1 h. DNA was purified by phenol-chloroform extraction, followed by ethanol precipitation (2 mg/mL glycogen, 500 μL 100% ethanol, 15 min on ice). Pellets were washed with 100% ethanol and resuspended in 30 μL 0.1x TE buffer.

### Next Generation Sequencing

ChIP and CUT&RUN DNA was quantified using the Qubit dsDNA HS Assay Kit (Invitrogen, Cat. No. Q32851). Libraries were prepared with the NEBNext Ultra II DNA Library Prep Kit (NEB, Cat. No. E7645S), following the standard protocol for ChIP and a modified protocol for CUT&RUN (39). Sequencing (2 x 60□bp paired-end) was performed on a NextSeq 2000 (LAFUGA, Gene Center Munich, Germany), generating ∼20 million reads per ChIP sample and ∼5 million per CUT&RUN sample.

### Data Analysis

For CUT&RUN, adapter sequences at the end of the reads were trimmed by the Cutadapt software (40). Then, the reads were aligned to the *Drosophila melanogaster* genome (dm6) by using Bowtie2 (41) with the parameters *-end-to-end --very-sensitive --no-mixed --no-discordant -I 10 -X 700*. SAMtools (42) and BEDTools (43) were utilized to convert the generated SAM files to BAM and BED files, respectively. BigWig coverage files were generated using the deepTools (44) bamCompare command with the parameters *--scaleFactorsMethod readCount --operation ratio --binSize 10 -- smoothLength 50* and normalized to pre-immune controls. The peaks were called by MACS2 in paired-end mode, with a fold change over the control of 4 and a q-value of 0.01. For ChIP data, the same pre-processing steps were followed, using the following bowtie2 alignment parameters: *-p 24 -- end-to-end --very-sensitive --no-unal --no-mixed --no-discordant -I 0 -X 400*, and with a fold change of 6 for MACS2 peak calling. Motif analysis was performed on MEME Suite (45).

Downstream analysis after peak calling was performed in the R environment (46). Overlapping peaks among the replicates were determined using the ChIPpeakAnno (47) package and annotated to genomic features by the ChIPSeeker (48) package. GO terms were interpreted with the clusterProfiler (49). Summary heatmaps were generated with ComplexHeatmap (50). Heatmaps at TSS and peak centers were plotted by using deepTools (44).

## Results

### Verification of Xbp1-DOM-A interaction

The transcription factor Xbp1 was identified in an IP-MS analysis conducted under stringent conditions, along with the other subunits of the DOM-A complex (3). The tight association suggested a functional relationship between Xbp1 and the epigenetic regulator. To verify this interaction and enable subsequent functional studies, we raised antibodies against the full-length recombinant Xbp1(transcript variant RE) in guinea pig and rabbit. Towards this end, recombinant Xbp1 was expressed in bacteria, purified by Ni-NTA affinity chromatography (Supplementary Figure S1A, B), and used as an antigen. The protein was expressed as a single species with a molecular weight of around 80 KDa.

The specificity of the antisera was validated in *Drosophila* S2 cells. The rabbit polyclonal antibodies were used to immunoprecipitate Xbp1, while the guinea pig antiserum detected the protein by immunoblotting (see antibodies in Supplementary Table 1). We probed whole cell lysates of either untreated cells or cells in which *xbp1* expression had been reduced by RNA interference (RNAi, see sequences in Supplementary Table 2). An RNAi control directed at irrelevant sequences of glutathione-S-transferase (*gst*) of *Schistosoma japonicum* served to monitor non-specific effects.

An early study had shown that Xbp1 is expressed at very low levels in unstressed cells and a doublet of bands becomes detectable upon induction of the UPR by DTT treatment (27). Accordingly, we detected only a very faint doublet of bands around the expected size range in untreated proliferating cells (Figure 1A). The intensity of the signal increased upon non-specific RNAi-treatment, suggesting an elevated level of stress, but was completely abolished by RNAi against *xbp1*. Signals from cross-reacting (presumably cytoplasmic) proteins were not affected. Immunoprecipitation strongly enriched a doublet of bands, likely corresponding to two different isoforms (27) (Figure 1B),

**Figure 1.**
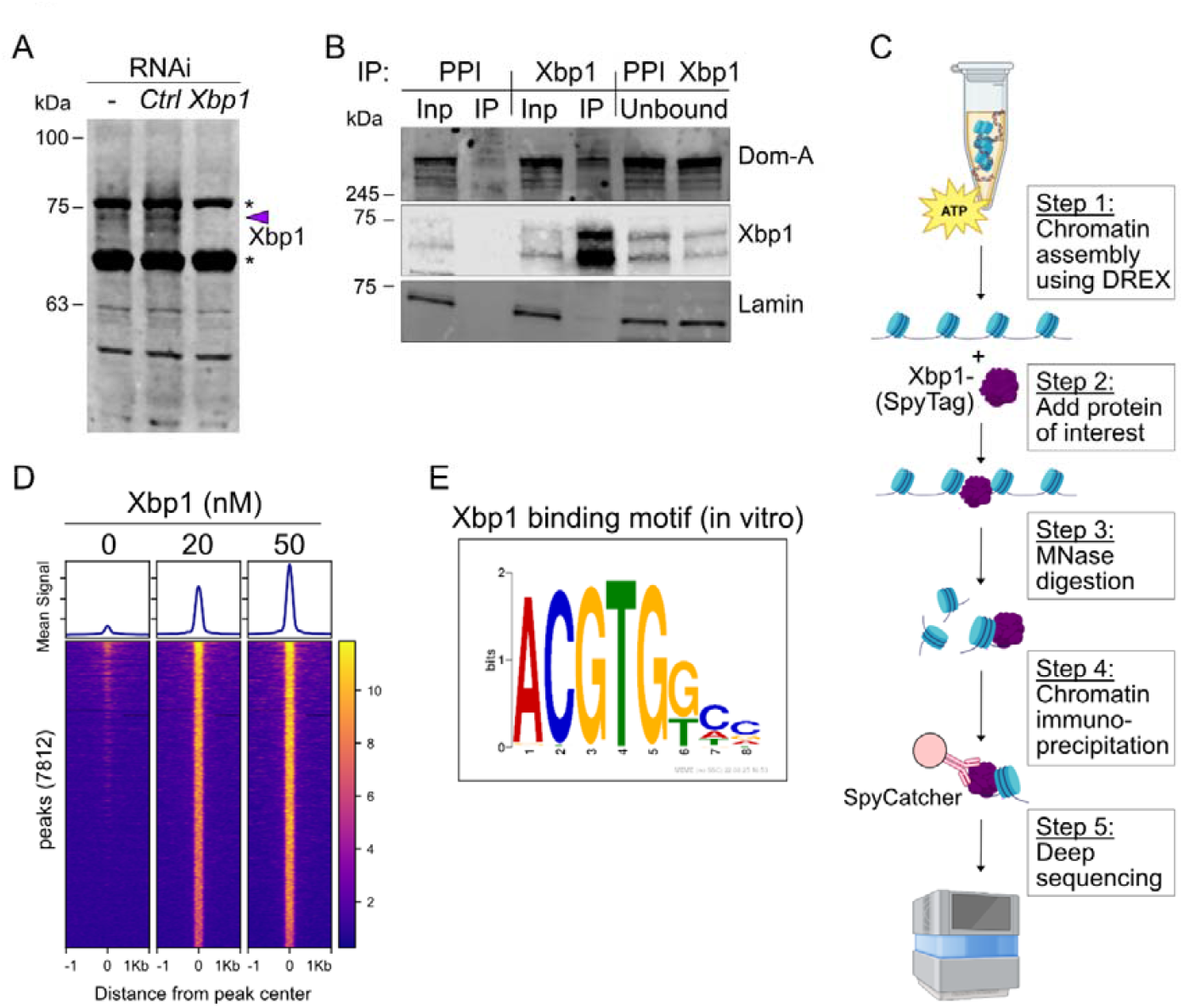
Characterization of Xbp1-Dom-A interaction and DNA binding motif. (A) Immunoblot analysis of Xbp1 expression in S2 cells. Whole cell extracts from untreated cells, cells treated with control RNAi (Ctrl), or Xbp1 RNAi were probed for Xbp1 (indicated by arrow) with a new Xbp1 antibody. The band corresponding to Xbp1 is indicated by the arrow; molecular weight markers are depicted on the left. Non-specific cytoplasmic proteins are marked with asterisks. (B) Co-immunoprecipitation (Co-IP) of Xbp1 and Dom-A. Xbp1 was immunoprecipitated from nuclear lysates from S2 cells with the pre-immune serum (PPI) as specificity control. Ten percent of input (Inp), precipitate (IP), and unbound fractions were analyzed by immunoblotting with antibodies for Dom-A, Xbp1, and lamin (as loading control). (C) Overview of the strategy used to identify Xbp1 binding sites in a chromatin-reconstituted *Drosophila* genome. Extracts from preblastoderm *Drosophila* embryos were used to assemble complex chromatin on genomic DNA. Recombinant Xbp1-SpyTag was then incubated with the reconstituted chromatin and crosslinked with formaldehyde. The chromatin was subsequently fragmented using MNase, and Xbp1-bound complexes were isolated via the SpyTag/SpyCatcher2 system. (D) Addition of 20 or 50 nM of Xbp1 to chromatin-reconstituted genomic DNA identified 7812 peaks at 50 nM Xbp1 from the average of 2 replicates. The heatmap and cumulative plot of Xbp1 ChIP signal in a 2 Kb window across these binding sites are shown. To generate the coverage signal, profiles were normalized to the input. (E) MEME motif analysis identified the displayed motif that is most strongly enriched among the Xbp1 binding sites.

The interaction between Dom-A and Xbp1 had originally been detected by Dom-A IP-MS. We validated this result through a reciprocal experiment, in which we immunoprecipitated Xbp1 from an S2 nuclear lysate and monitored associated Dom-A by immunoblotting. Indeed, Dom-A was retrieved upon Xbp1 IP, but not with the preimmune serum control (Figure 1B).

### Identification of the Xbp1 recognition sequence motif in chromatin-reconstituted fly genome

To our knowledge, Xbp1 binding sites in *Drosophila* have not been analyzed genome-wide, and a binding motif is unknown. To determine the Xbp1 sequence interaction motif, we monitored the binding of recombinant to genomic binding sites in reconstituted chromatin *in vitro* (Figure 1C). In short, physiological chromatin was assembled on purified genomic DNA in a cell-free chromatin reconstitution system derived from preblastoderm *Drosophila* embryos (51, 52). Xbp1 was allowed to bind to the complex chromatin substrate, interactions were fixed by formaldehyde crosslinking, and the chromatin was fragmented by digestion with Micrococcal Nuclease (Supplemental Figure S1C). Xbp1 was immunoprecipitated using SpyCatcher and the associated DNA subjected to paired-end sequencing. As a quality control for the chromatin reconstitution, we mapped phased nucleosomes around 2237 binding sites for su(Hw) [Suppressor-of-hairy-wing], an insulator complex endogenous to the embryonic extract (Supplemental Figure S1D) (51).

Recombinant Xbp1 interacted robustly with thousands of sequences in the fly genome in a dose-dependent manner. At a concentration of 50 nM Xbp1, 7812 peaks were called (Figure 1D). Motif enrichment analysis identified the ‘ACGTG’ core sequence motif in 95% of the peaks (Figure 1E), which is compatible with the known binding site for the human hXBP1 (53).

### Xbp1 associates with promoters controlling ER function, UPR, and protein trafficking

Xbp1 expression is regulated post-transcriptionally through an unconventional, cytoplasmic splicing by IRE1 during the UPR. The removal of a short 23-nucleotide intron leads to a frameshift in *Xbp1* mRNA, which is translated into the functional Xbp1 transcription factor (Figure 2A). The UPR can be induced experimentally by the addition of the reducing agent dithiothreitol (DTT) to the cell medium (27). The spliced mRNA is detected by PCR within 30 minutes of adding 1 or 5 mM DTT to the medium of S2 cells (Supplementary Figure S2A). A basal level of spliced mRNA isoform is already detectable in the absence of DTT, which may be due to a mild stress under our growth conditions or represent gene regulatory functions in physiological ER homeostasis (29). The level of *Xbp1* mRNA was only slightly increased upon DTT treatment, suggesting that transcription of the *Xbp1* gene is not the primary regulatory principle in the UPR. In contrast, the mRNA levels encoding the ER-resident chaperone Hsc70-3 (dBip) were induced up to 8-fold after DTT treatment (Supplementary Figure S2B). In the subsequent nucleosome mapping experiments, we treated the cells with 5 mM DTT for 1 hour, which robustly increased the levels of Xbp1 (Figure 2B). Of note, this enhanced signal is completely abolished by RNAi against Xbp1, once more confirming the specificity of the antibody employed, as well as the effectiveness of our depletion strategy.

**Figure 2.**
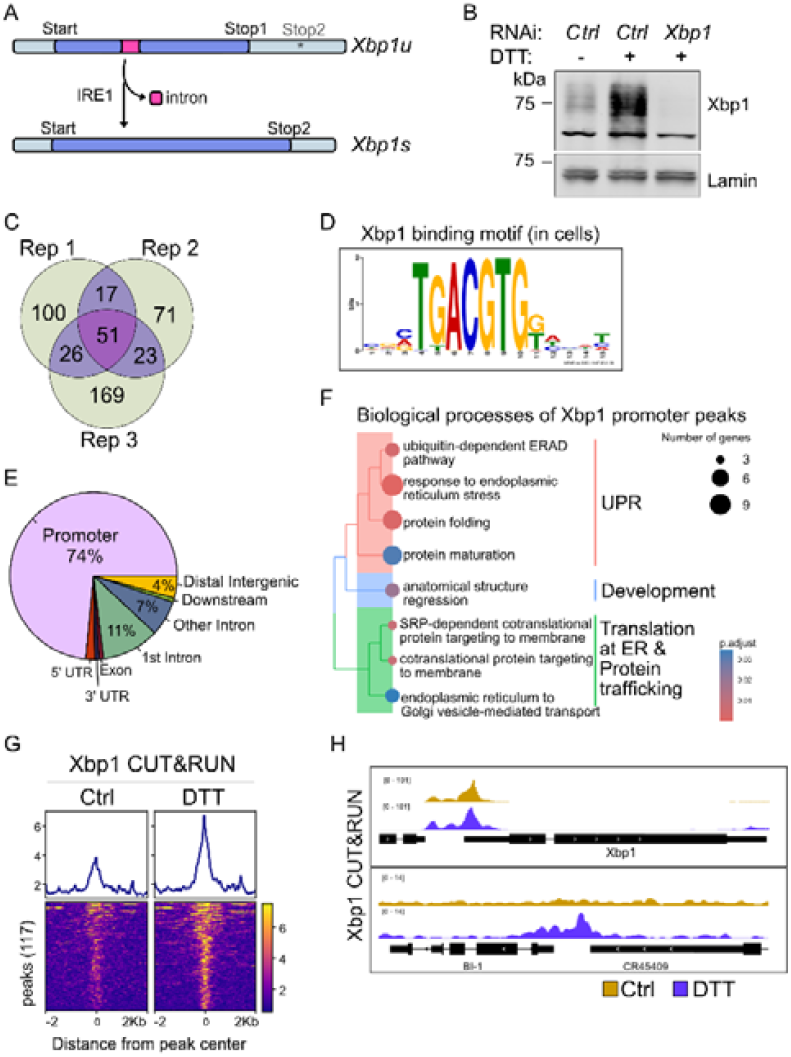
CUT&RUN identifies Xbp1 targets regulating ER function. (A) Illustration of unconventional splicing of *Xbp1* mRNA by IRE1 during the UPR. *Xbp1u*: unspliced *Xbp1* mRNA, *Xbp1s*: spliced form. For details see text. (B) Xbp1 expression increases during activation of the UPR upon DTT treatment of S2 cells. Nuclear lysates from cells subjected to Xbp1 or control RNAi, and treated or not with 5 mM DTT, were analyzed by immunoblotting using antibodies against Xbp1 and the loading control lamin. (C) Identification of high-confidence Xbp1 binding sites. CUT&RUN was performed with anti-Xbp1 antibodies in DTT-treated S2 cells. Peaks were called in each of three biological replicates. The Venn diagram identifies the overlapping sites. Peaks called in at least two out of three replicates (n=117) were considered high-confidence binding events for downstream analysis. (D) MEME motif enrichment analysis of *in vivo* Xbp1 binding sites from (C). (E) Genomic feature annotation of 117 high-confidence Xbp1 binding sites. The pie chart displays the relative distribution of peaks across the indicated genomic features. (F) Gene Ontology (GO) term analysis of Xbp1 binding sites. Promoters identified in (E) were subjected to Biological Process (BP) analysis to define significant GO terms represented in the dendrogram. Similar terms were clustered and three main categories were highlighted to the right. The node size indicates the number of genes in each term and p-values are color-coded. (G) The heatmap and cumulative plot of Xbp1 CUT&RUN signal in a 4 Kb window across the high-confidence binding sites are shown. Xbp1 coverage signals were normalized to their respective PPI controls for each replicate, and then three biological replicates were averaged. DTT-treated and untreated samples are shown. (H) Genome browser snapshots of the Xbp1 CUT&RUN coverage at promoters of *Xbp1* and *BI-1* genes are shown in the absence (Ctrl-yellow) or presence (DTT-purple) of DTT.

To map the genomic binding sites for Xbp1, we conducted CUT&RUN (54) in DTT-treated S2 cells along with the untreated control. Three biological replicates were analyzed separately and those peaks called in at least two replicates were pooled, resulting in 117 binding sites detected in DTT-treated cells that we consider robust binding events (Figure 2C). The majority of these sites map to promoter regions (Figure 2E). A MEME motif enrichment analysis identified “TGACGTG” as the most highly enriched motif. It is present at least once in 58 of the 117 sites (Figure 2D). While the identification of this motif fits our binding site determination *in vitro*, many robust *in vivo* binding sites of Xbp1 do not contain such a motif. Interestingly, the *in vivo* motif contains a 5’ TG di-nucleotide extension to the core “ACGTG” sequence identified in the genome-wide *in vitro* ChIP, which is similar to the motif of human Xbp1 (53)(Figure 1D). Interestingly, when we searched for known motifs related to the Xbp1 “TGACGTG” motif, we observed an overlap with the consensus sequence of another UPR effector, Atf6 (Supplementary Figure S2C).

Functional categorization of promoters bound by Xbp1 reveals that most associated genes participate in maintaining proper ER function, the UPR, and protein trafficking (Figure 2F). The robust binding of Xbp1 to the 117 sites upon DTT treatment is illustrated in Figure 2G as increasing CUT&RUN signal intensities in Figure 2G. Xbp1 binding at some of these sites is detectable already in the absence of DTT, as expected from its baseline expression. Interestingly, the sites differ in their occupancy under uninduced conditions. For example, Xbp1 already binds strongly to its own promoter in the absence of DTT, and this binding is not much enhanced by DTT (Figure 2H). On the other hand, low-level binding at the promoter of the *bax inhibitor 1(bI-1)* gene, encoding an ER membrane protein involved in the UPR, is strongly increased upon DTT treatment, illustrating the role of Xbp1 in promoting the UPR. Further examples for either constitutive or DTT-induced binding are shown in Supplementary Figure S2D.

### DOM-A binds to a diverse set of promoters, but only few contain Xbp1 binding sites

Since the genomic binding sites of Dom-A had not been determined before, we once more employed CUT&RUN. Signals that were called in two out of three biological replicates (n = 1324) were considered robust binding sites and analyzed further (Figure 3A). Evidently, Dom-A binds to many more sites than Xbp1, presumably because it is involved in processes other than ER homeostasis and the UPR as well.

**Figure 3.**
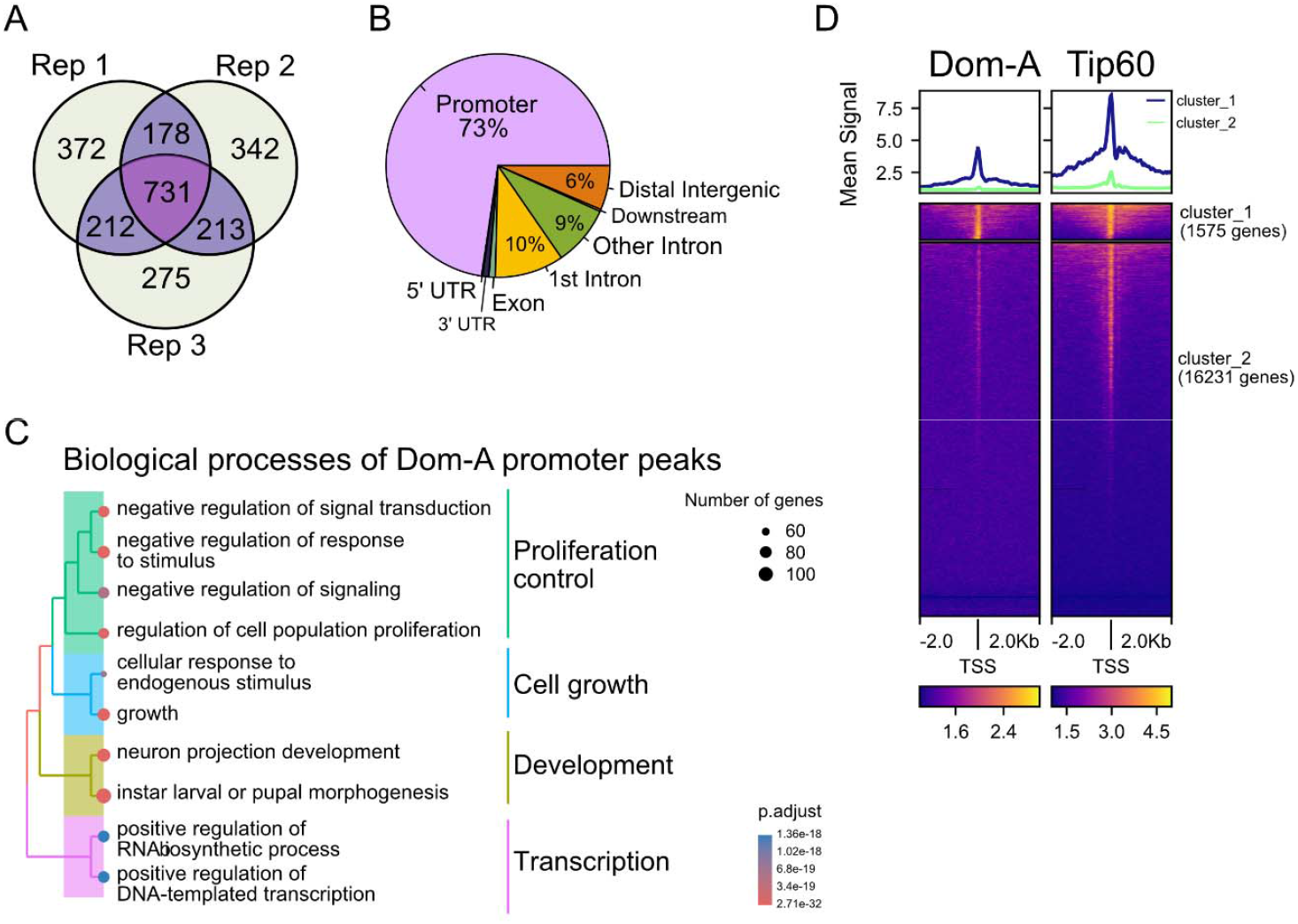
Dom-A binds to a variety of gene promoters and overlaps with Tip60 binding sites. (A) Venn diagram demonstrating the number of peaks in three biological replicates of Dom-A CUT&RUN in S2 cells. Data were normalized to the IgG control for peak calling. Overlapping peaks were identified among the replicates, and the peaks called in two or more replicates (n=1324) were considered high-confidence binding events and included in subsequent analyses. (B) Genomic feature annotation of high-confidence Dom-A binding sites. The pie chart displays the percentage distribution of peaks across genomic features. (C) Dendrogram representing a GO-term tree plot of Dom-A target genes. The binding sites located in gene promoters (from B) were analyzed for Biological Process (BP) enrichment. Significant GO-terms were further clustered into 4 main categories, indicated on the right. The number of genes in each term and their p-values are denoted by the node size and color scale, respectively. (D) Heatmap and cumulative plot showing Dom-A and Tip60 enrichment around 17806 transcription start sites (TSS) in a window of 4 Kb. The signals represent the average of three biological replicates, normalized to the corresponding IgG controls. Heatmap is clustered by k-means = 2 based on Dom-A.

Once more, the majority of Dom-A binds at promoter regions (Figure 3B). The functional categorization of the corresponding genes highlights the broad terms ‘proliferation control’, ‘cell growth’, ‘development’ and ‘transcriptional control’ (Figure 3C). This is in line with our recent findings concerning the DOM-A subunit Tip60, that led us to conclude about a major role for the DOM-A complex in cell growth and proliferation. Comparison of the published Tip60 CUT&RUN profiles with the current Dom-A promoter binding maps shows a high degree of overlap (Figure 3D), confirming the earlier conclusion that Dom-A and Tip60 are two effector enzymes of the same complex.

### Xbp1 recruits Dom-A to promoters of genes involved in ER homeostasis and UPR

The availability of CUT&RUN profiles for Xbp1 and Dom-A allows us to determine the degree of their colocalization in the genome. In support of our hypothesis that the strong interaction of Xbp1 with the DOM-A complex may serve to target DOM-A to Xbp1-regulated promoters, we found Dom-A robustly localized at the 117 high-confidence Xbp1 binding sites (Figure 4A).

**Figure 4.**
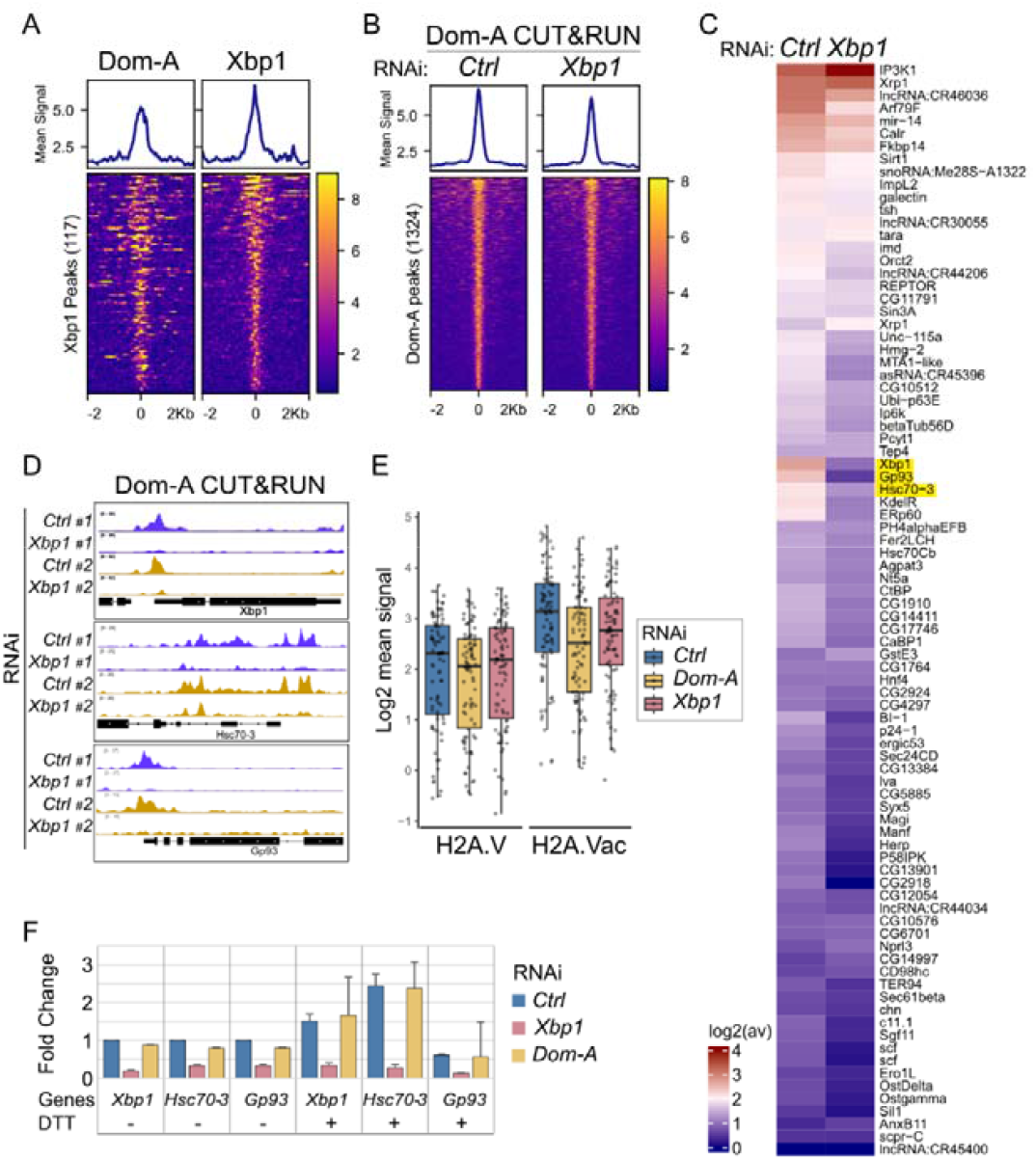
Xbp1 targets Dom-A to ER homeostasis and UPR genes. (A) Heatmap and cumulative plot of CUT&RUN reads comparing the enrichment of Dom-A and Xbp1 in a 4 Kb window centered at high-confidence, DTT-induced Xbp1 binding sites. Signal sorted based on decreasing Xbp1 intensities. (B) Heatmap and cumulative plot of Dom-A CUT&RUN signals in a 4 Kb window centered at high-confidence Dom-A binding sites in control (Ctrl) S2 cells and upon Xbp1 depletion. Profiles from 3 biological replicates were each normalized to their respective IgG controls and then averaged. (C) Heatmap of Dom-A CUT&RUN signal from (B) at high-confidence Xbp1 target promoters. Average signal intensities were calculated for the promoters within a 1 Kb window centered on the TSS (± 500 bp) and log2-transformed. Gene symbols for the corresponding transcripts are depicted to the right. The three loci that exhibit the most pronounced decrease in Dom-A binding are highlighted. (D) Genome browser snapshots of Dom-A coverage in control cells and upon Xbp1. Two biological replicates show the three loci highlighted in (C) (*Xbp1, Hsc70-3* and *Gp93*). (E) Box plot representation of the average H2A.V and H2A.Vac signal at Xbp1 target promoters (TSS ± 500 bp) upon depletion of Dom-A or Xbp1, relative to non-targeting RNAi (Ctrl). Each dot corresponds to a TSS. Mean signals from 3 replicates in a 1 Kb window centered around TSS was log2-transformed. (F) RT-qPCR analysis of *Xbp1, Hsc70-3* and *Gp93* mRNA levels. Fold changes relative to the RNAi control of the corresponding genes were shown for both Xbp1- and Dom-A-depleted cells.

To test the hypothesis that Xbp1 may target Dom-A to sites of action, we monitored the Dom-A chromatin binding upon Xbp1 depletion. The profile of Dom-A binding to its 1324 high-confidence sites shows that, overall, Xbp1 depletion has little effect on Dom-A occupancy (Figure 4B). However, the cumulative signal across these sites indicates reduced Dom-A binding at a subset of locations (Figure 4B, Supplementary Figures S3A, B). Examining the promoter regions of the 117 high-confidence Xbp1 binding sites, we observed that Dom-A binding was reduced to varying degrees in the absence of Xbp1 (Figure 4C). As inferred from the functional categorization in Figure 1F, the majority of these promoters regulate genes involved in ER homeostasis and the UPR. The three most affected genes are highlighted in the list to the right of Figure 4C. These are the *Xbp1* gene itself and genes encoding two prominent chaperones involved in the UPR, *Hsc70-3* and *Gp93*. Genome browser snapshots show that these promoters are strongly bound by Xbp1 and the depletion of the TF leads to a strong reduction of Dom-A in all three cases (Figure 4D).

The DOM-A complex combines two enzymatic activities, the H2A.V exchange ATPase Domino-A and the acetyltransferase Tip60. In line with this notion, depletion of Dom-A reduced the levels of H2A.V and of H2A.V acetylation at Xbp1 target promoters to varying degrees (Figure 4E). RNA interference with Xbp1 expression also led to reduced H2A.V acetylation, although to a lesser degree.

The importance of Xbp1 as a transcription regulator became clear when we measured the expression of the three model genes, *Xbp1, Hsc70-3* and *Gp93*, following Xbp1 depletion. Both basal and DTT-induced mRNA levels were markedly reduced in the absence of Xbp1 (Figure 4F). However, depletion of Dom-A did not diminish transcription (see discussion).

### Stabilization and ‘reverse recruitment’ of Xbp1 by Dom-A

The recruitment of Dom-A to a relatively small number of high-confidence Xbp1 target genes seems at odds with our initial finding that Xbp1 was identified as a detergent-resistant interactor of the DOM-A complex in an IP-MS screen. Conceivably, Xbp1-Dom-A interactions may not be limited to the strongest Xbp1 binding sites. To explore this, we examined potential Xbp1 binding across all 1324 high-confidence Dom-A binding sites. Interestingly, we detected a distinct signal that mirrors Dom-A binding (Figure 5A). Considering that the vast majority of the Dom-A sites do not contain an Xbp1 recognition sequence motif, we speculate that Xbp1 may be localized at these sites due to its interaction with DOM-A. In such a scenario, DOM-A would recruit Xbp1, rather than the other way around. To test this idea, we depleted Dom-A and monitored the genomic binding of Xbp1. To our surprise, we found the Xbp1 binding to Dom-A sites drastically reduced (Figure 5A). This general observation is illustrated by visualizing the binding of Xbp1 to its key target genes, *Xbp1, Hsc70-3* and *Gp93* (Figure 5B). Remarkably, Xbp1 binds much less to all three promoter regions, if Dom-A is depleted.

**Figure 5.**
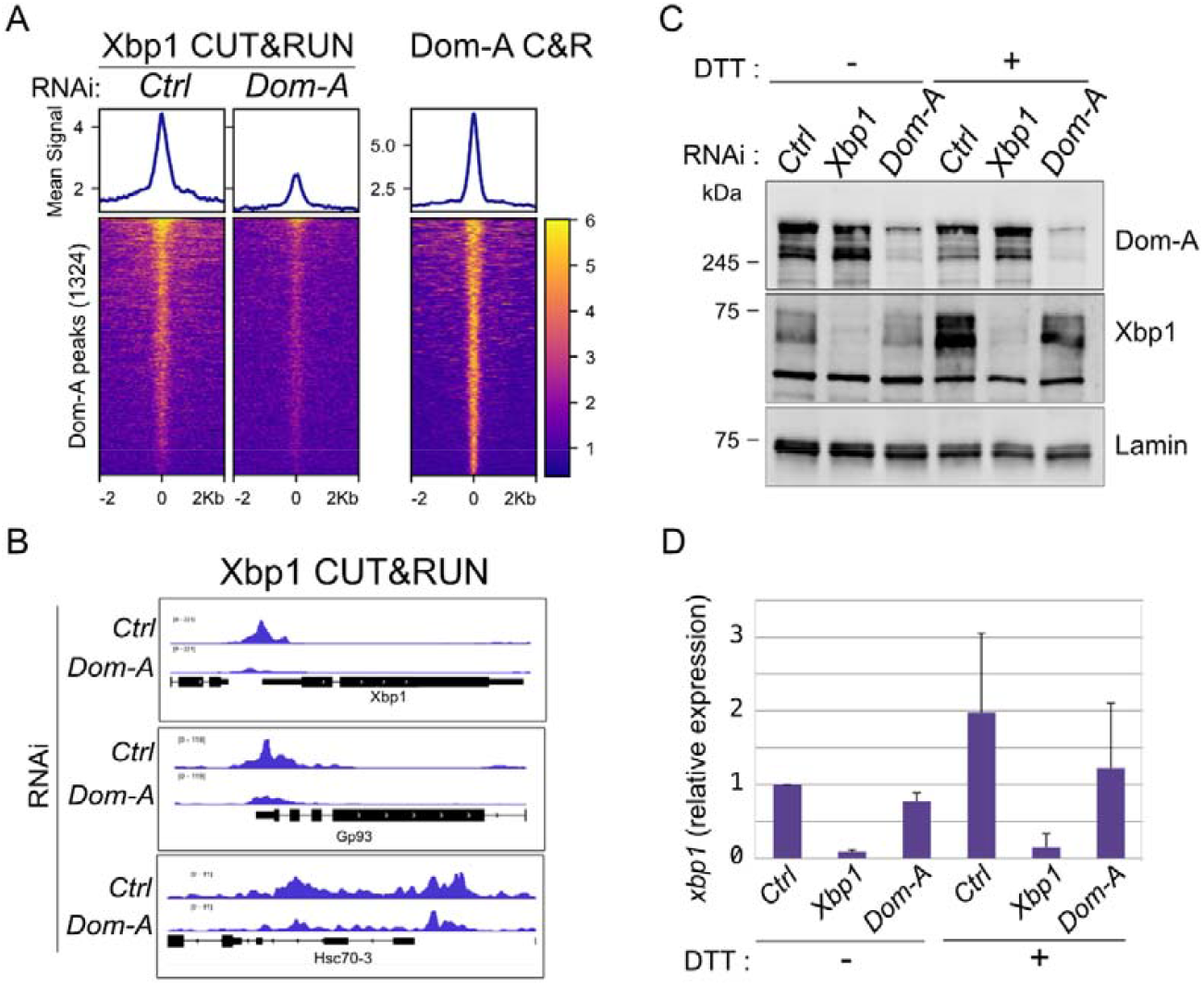
Dom-A is required for Xbp1 stabilization and chromatin interactions. (A) Heatmap and cumulative plot of Xbp1 (left panels) and Dom-A (right panels) CUT&RUN signals in a 4 Kb window centered at 1324 high-confidence Dom-A binding sites in S2 cells. The Xbp1 binding was monitored in control cells and upon Dom-A depletion as indicated (RNAi). Profiles from 3 biological replicates were each normalized to their respective IgG controls and then averaged. Signal sorted based on decreasing intensities. (B) Genome browser snapshot of Xbp1 coverage in control cells (Ctrl) or upon Dom-A depletion at promoters of *Xbp1, Gp93* and *Hsc70-3* genes. (C) Immunoblot analysis of Dom-A and Xbp1 protein levels after reciprocal factor depletions. Nuclear lysates were prepared from S2 cells treated with RNAi against Xbp1 or Dom-A, or with a non-specific RNAi control (Ctrl) with or without DTT treatment. Blots were probed for Dom-A, Xbp1, or lamin (loading control), as indicated. (D) Quantification of Xbp1 protein levels from (D). Fold changes over control RNAi without DTT were calculated after normalization to the lamin signal. Error bars represent the standard deviation from two biological replicates.

Reduced chromatin binding could result from lower protein abundance. Immunoblot analysis showed that RNAi directed against Xbp1 markedly decreased Xbp1 protein levels, as expected, while Dom-A levels remained unchanged (Figure 5C, Supplementary Figure S3C). Thus, the substantially reduced Dom-A binding at Xbp1 target promoters (Figures 4B, C) cannot be attributed to decreased Dom-A protein levels. In contrast, depletion of Dom-A reduced the Xbp1 levels by roughly 50% (Figures 5C, D, Supplementary Figure S3C). Conceivably, the interaction between Xbp1 and DOM-A contributes to the stabilization of the TF, analogous to the stabilization of Tip60 within the DOM-A complex (3). Depletion of Tip60 did not affect Xbp1 protein levels (Supplementary Figure S3C), unlike mammalian Xbp1, whose stability has been shown to be affected by P300 acetylation (55, 56).

## Discussion

### Intersection of Xbp1 and DOM-A functions in ER quality control and cell cycle progression

The atypical, detergent- and benzonase-resistant association of Xbp1 with the Dom-A complex suggests a functional interaction of the transcription factor (TF) with the epigenetic coregulator (3). Xbp1 is a critical regulator of ER proteostasis. If the level of unfolded proteins exceeds the folding capacity of the ER, cells mount the Unfolded Protein Response (UPR), which involves increasing the folding and secretory capacity of ER, dampening translation and ribosome biogenesis and enhancing the degradation of misfolded proteins (57). Xbp1, along with the transcription factor ATF6, orchestrates the transcriptional arm of the UPR by boosting the transcription of genes encoding chaperones and other factors that improve the folding and secretory capacity of the ER (36).

Mammalian XBP1 and ATF6 can form homo-or heterodimers that bind to sequence motifs in the promoters of UPR target genes (58). The general concept of TF-mediated targeting of an epigenetic coregulator to a promoter involves binding of a TF to a small sequence motif and interaction of the coregulator with the DNA-bound TF. *Drosophila* Xbp1 is poorly studied so far. We experimentally identified the Xbp1 core recognition motif through genome-wide biochemical mapping of binding sites in the context of *in vitro-reconstituted* physiological chromatin (59). The sequence motif is consistent with the known binding site for human hXBP, reflecting the evolutionary conservation of XBP factors (53, 60). Because the reconstituted chromatin resembles dynamic embryonic chromatin characterized by high nucleosome mobility, the assay identifies all possible binding sites. Not surprisingly, many of those sequences are not bound in the mature cellular chromatin, where productive binding usually requires cooperativity with other TFs and accessible sites (61).

Xbp1 and Atf6 cooperate and complement each other, but may also establish some functional redundancy (57). In line with such functional interactions, we found that the Xbp1 recognition motifs overlap with the known binding motif of *Drosophila* Atf6.

An involvement of the DOM-A complex in fine-tuning the UPR is well in line with our knowledge about this global regulator. We previously observed that DOM-A, through its effector module Tip60, controls many aspects of cell growth and cell cycle progression (7). DOM-A binds to thousands of promoters, where Tip60 acetylates the histone variant H2A.V (H2A.Z in mammals) at transcription start sites. Depletion of Tip60 leads to reduced H2A.V acetylation and impaired transcription of many genes encoding critical proteins for energy provision, biosynthesis pathways, cell cycle progression and division. In addition to its role in transcription, Tip60 acetylates numerous proteins involved in all aspects of cell growth and proliferation, including nucleolar proteins and ribosome subunits (7). Because smooth progression through the cell cycle requires properly folded proteins, DOM-A may be well suited to integrate proteostasis signals with general regulation of cell growth and proliferation.

### Xbp1 recruits Dom-A to a small subset of DOM-A target promoters

The localization of Dom-A/Tip60 at thousands of promoters may, to some extent, involve recognition of general features of active promoters by reader domains in DOM-A subunits, such as acetylated nucleosomes, or extended nucleosome-free regions (16, 18). In addition, many studies of the orthologous yeast NuA4 and human P400 complexes provide examples for their recruitment by transcription factors (TFs) (62). Accordingly, it is likely that TFs contact DOM-A via the Nipped-A subunit (Tra1 in yeast, TRRAP in mammals). This conserved, large subunit of DOM-A (3) provides a central, if not the only, surface for TF interactions (62). Mapping the physical requirements underlying the Xbp1/ Nipped-A interactions remains an interesting future task.

In line with the idea of multiple targeting principles for DOM-A, including other TFs, Xbp1 mapped to a small subset of Dom-A-bound promoters. The high-confidence Xbp1 promoter binding sites identified a set of genes highly enriched for proteins involved in ER proteostasis. Curiously, only about half of these robust binding sites contain an Xbp1 recognition motif. Focused analyses on key genes of the UPR, such as *Xbp1, Hsc70-3*, and *Gp93*, confirmed that these promoters fulfil the criteria for a functional recruitment of DOM-A by Xbp1: depletion of Xbp1 reduced their transcription, abolished Dom-A binding, and lowered levels of acetylated H2A.V. However, we did not observe a reduction of transcription upon Dom-A depletion. This may be due to coactivator redundancy: the Nipped-A subunit, the likely target of Xbp1, is also a subunit of the SAGA acetyltransferase complex, a well-known, strong activator of transcription (62). The recruitment of SAGA through Nipped-A was shown earlier for the transcription factor Clk (63). Xbp1 may thus target both SAGA and DOM-A via a similar mechanism. A failure to document a specific reduction of transcription may also be due to a technical limitation: Dom-A depletion leads to lowered transcription of many genes, including housekeeping factors, such as GAPDH and 7SK, which we commonly use for normalization. A specific effect above a background of general transcription repression is difficult to document.

Interestingly, Xbp1 appears to be involved in a positive feedback loop activating its own promoter, which contains Xbp1 motifs and shows Dom-A recruitment. Of note, the expression of the Xbp1 gene is not only regulated at the level of transcription, but also by an unconventional processing of the primary transcript to yield functional *Xbp1* mRNA. High levels of misfolded proteins will thus lead to an increase of functional *Xbp1* mRNA and protein, which will boost the transcription of its own gene.

### Complex function interactions between Xbp1 and Dom-A

For critical genes of the UPR, such as *Xbp1, Hsc70-3* and *Gp93*, we found that Xbp1 is required for Dom-A binding, in line with a classical recruitment paradigm. Remarkably, we also observed the converse, namely that depletion of Dom-A led to strongly reduced levels of promoter-bound Xbp1. This phenomenon cannot be explained by reduced transcription of the *Xbp1* gene in the context of a feedback loop since *Xbp1* mRNA levels are unaffected in the absence of Dom-A. Depletion of Dom-A leads to overall reduced Xbp1 levels by 50% or more, suggesting that the interaction of the TF with the DOM-A complex stabilizes the protein. Whether the lower amounts of Xbp1 are sufficient to explain its absence at some of its strongest binding sites is presently unclear.

The robust localization of Xbp1 to sites containing consensus binding motifs is presumably due to direct DNA binding. However, we also score much fainter Xbp1 interactions at hundreds of high-confidence Dom-A binding sites, most of which lack an Xbp1 consensus motif. These Xbp1 binding events are strongly reduced upon Dom-A depletion, suggesting some kind ‘reverse targeting’, where Dom-A is attracted by other principles and Xbp1 somehow piggybacks on Dom-A.

We can only speculate about the functional implications of such interactions. Many transcription factors are found associated with sites in chromatin that lack known sequence recognition motifs and they may have context-specific, opposing functions as activators or repressors (64). We concluded earlier that the DOM-A complex with its Tip60 effector is a central facilitator of the proliferative state of cells (7). Xbp1 has the opposite function, since the UPR slows down protein synthesis and cell proliferation to give the cell time to resolve the stress situation. We imagine complex physical and functional interactions between Xbp1 and DOM-A, involving binding of Xbp1 to the DOM-A complex (and hence its stabilization), recruitment of DOM-A to a small set of UPR-related promoters via direct Xbp1-DNA interactions and sequence-independent localization of Xbp1 to hundreds of Dom-A target sites through DOM-A interactions. Such interactions theoretically bear potential to allosterically modulate the functions of chromatin-bound DOM-A to negotiate the proliferative cues in the presence of unfolded proteins. Because many cells express a basal level of Xbp1 to assure ER proteostasis (36) a corollary of such a ‘moonlighting’ function of Xbp1 would be that DOM-A complexes are rarely fully active, but subject to constant regulation in accordance with requirements for growth and proliferation, such as proteostasis, energy availability, and DNA repair.

## Acknowledgements

We thank Silke Krause and Aline Campos Sparr for expert technical assistance, Carla Margulies for the critical discussions, and members of the Becker Lab for valuable input and critical reading of the manuscript. We thank T. Straub from the BMC Bioinformatics Unit for granting us access to the computational cluster, providing us with advice on bioinformatics and helpful scripts. We thank S. Krebs of LMU LAFUGA for Illumina sequencing. The graphical abstract was created with BioRender.

## Author contributions

GK: Conceptualization, Investigation, Data curation, Formal analysis, Validation, Writing—original draft, Writing—review & editing; PPB: Conceptualization, Funding acquisition, Supervision, Validation, Writing—original draft, Writing—review & editing): ZA: Conceptualization, Data interpretation, Supervision, Validation, Writing—original draft, Writing—review & editing).

## Supplementary data

Supplementary Data are available at the journal’s site online.

## Data availability

Sequencing data were deposited in ArrayExpress under accession code E-MTAB-15796.

