## Supplementary for "Reciprocal targeting of the unfolded protein response regulator Xbp1 and the Dom-A nucleosome remodeler in *Drosophila*"

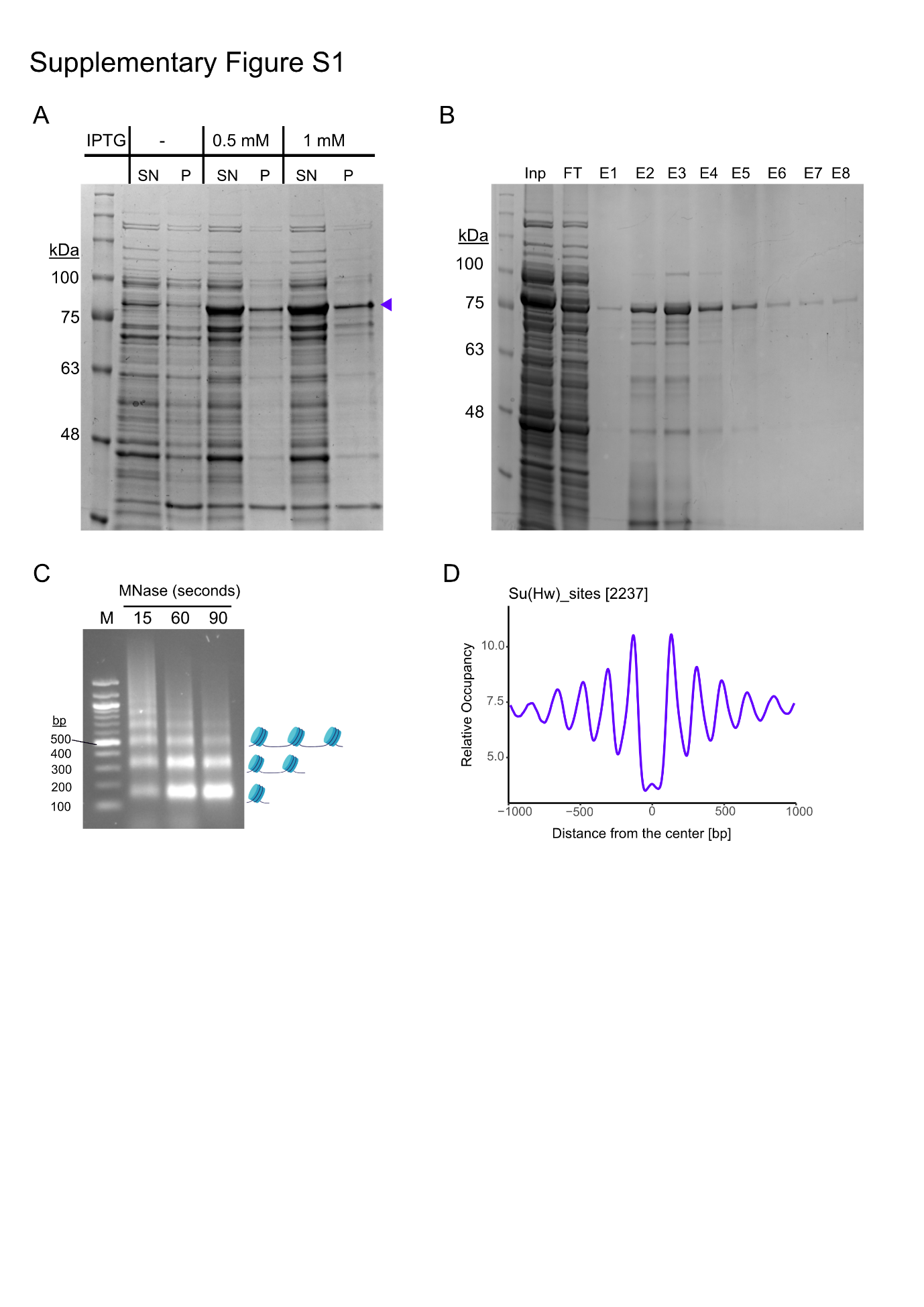

Supplementary Figure S1

(A) Bacterial expression of recombinant Xbp1 protein. Bacterial cultures with or without IPTG induction were fractionated to test the solubility of the protein (SN: supernatant, P: pellet). The Xbp1 band induced by IPTG is specified with the arrow above 75 kDa.
(B) Supernatant of the bacterial lysate was subjected to Ni-NTA beads to purify His-tagged Xbp1. Input (Inp), Flow Through (FT), and Elution (E1-E8) fractions resolved on a 10% SDS Gel.
(C) MNase digestion of in vitro-reconstituted chromatin. Following chromatin reconstitution, samples were treated with MNase for the indicated durations. The resulting DNA was purified and analyzed on an agarose gel, revealing the characteristic nucleosomal DNA ladder. M: Marker (NEB 100 bp). Mono-, di-, tri-nucleosomal DNA are depicted on the right.
(D) Nucleosome phasing at 2237 Su(Hw) binding sites in the Drosophila genome. Following MNase digestion, mononucleosomal DNA was purified and subjected to sequencing. Sequencing reads within a 2 kb window surrounding Su(Hw) sites were plotted as a cumulative profile, revealing nucleosome phasing patterns.

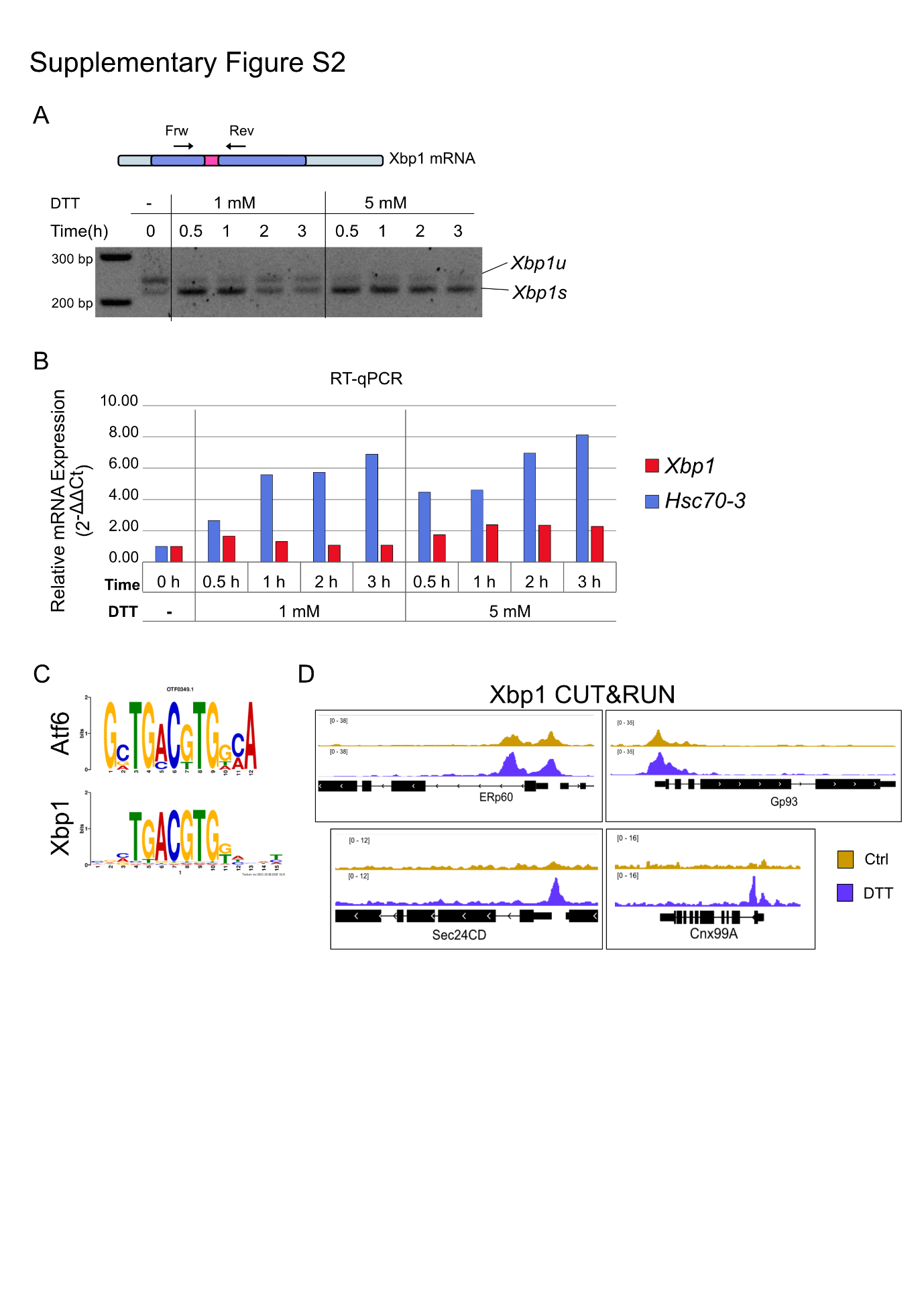

Supplementary Figure S2

(A) Agarose gel image showing the two forms of Xbp1 transcripts, the unspliced (Xbp1u) and the IRE1-spliced (Xbp1s). Cells were treated with 1 mM or 5 mM DTT for the indicated time points, after which total RNA was extracted and reverse-transcribed into cDNA. A primer set capable of amplifying both Xbp1 mRNA isoforms was used. The PCR products were resolved on a 3% agarose gel to distinguish between the unspliced (Xbp1u) and spliced (Xbp1s) forms, which differ by the removal of a 23-nt intron.
(B) RT-qPCR analysis of Xbp1 and Hsc70-3 (dBip) mRNA expressions upon DTT treatment of the samples in (A). Relative expressions were calculated by the ΔΔCt method. RpII140 was used as a reference gene for normalization.
(C) Tomtom motif similarity analysis matches the Xbp1 motif to the Atf6 motif.
(D) Genome browser views showing Xbp1 coverage signals at representative genes. Xbp1 enrichment over certain promoters (e.g., *ERp60* and *Gp93*) remains consistent regardless of stress induction, whereas at other promoters (e.g., *Sec24CD* and *Cnx99A*), enrichment increases upon DTT-induced stress.

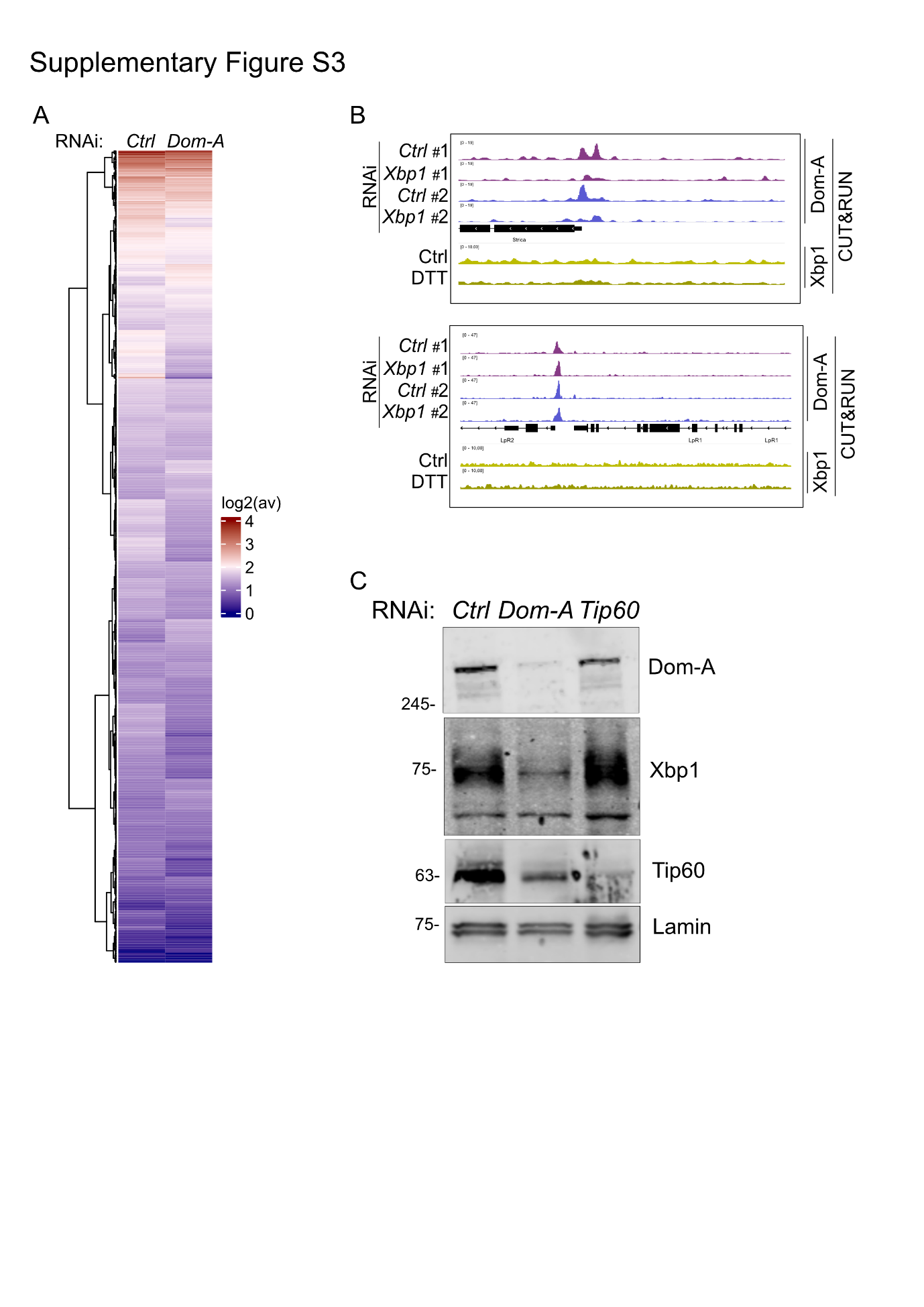

Supplementary Figure S3

(A) Summary heatmap showing Dom-A CUT&RUN signals at Dom-A target promoters following Xbp1 depletion compared to a non-specific control RNAi. The log_2_ average signal was calculated within a 1 kb window centered on the TSS of defined Dom-A target genes.
(B) Genome browser snapshots showing Dom-A and Xbp1 coverage across the indicated gene promoters. For Dom-A CUT&RUN, RNAi conditions are labeled on the left, with two biological replicates displayed in different colors. For Xbp1 CUT&RUN, the plotted signal represents the average of three biological replicates, with control and DTT-treated samples indicated on the left.
(C) Immunoblot analysis of Xbp1, Dom-A, and Tip60 upon Dom-A or Tip60 knockdown relative to non-specific RNAi control (Ctrl). Lamin served as a loading control. Molecular weights are shown on the left.

Table 1. Antibodies

| Antibody | Host Species | Dilution/Application | Source/Cat. No. | Reference |
| --- | --- | --- | --- | --- |
| a-Dom-A | Rat-monoclonal | 1:20/Western Blot | Peter B. Becker Lab/ 17F4 | Börner and Becker, 2016 |
| a-Dom-A | Rabbit-Polyclonal | 1:500/CUT&RUN | Peter B. Becker Lab/ SA8977 |  |
| a-Lamin | Mouse-monoclonal | 1:1000/Western Blot | Gift from H.Saumweber/  T40 |  |
| a-Tip60 | Rat-monoclonal | 1:20/Western Blot | Peter B. Becker Lab/ 11B10 | Scacchetti et al., 2020 |
| a-Xbp1 | Guinea Pig-Polyclonal | 1:1000/Western Blot | Peter B. Becker Lab/ SAC570 |  |
| a-Xbp1 | Rabbit-Polyclonal | 1:500/CUT&RUN | Peter B. Becker Lab/ SA1228 |  |
| a-H2A.V | Rabbit-Polyclonal | 1:150/CUT&RUN | Peter B. Becker Lab/ SA4871 | Börner and Becker, 2016 |
| a-H2A.Zac(K4,K7) | Rabbit-Monoclonal | 1:300/CUT&RUN | Cell Signaling/#75336S (D3V1I) |  |
| Human IgG1-FcSpyCatcher3 | Recombinant-Purified | 1:1000/ChIP | Bio-Rad/#TZC009 | Michael et al., 2023 |
| Normal Rabbit IgG | Purified | CUT&RUN | Cell Signaling/#2729 |  |

Table 2. Primers

| Primer | Sequence | Application |
| --- | --- | --- |
| RpII140_F | GCGCTATGGGTAAGCAAGCT | RT-qPCR |
| RpII140_R | TCACAAGTGGCTTCATCGGA | RT-qPCR |
| Hsc70-3_F | ATTACTGGCCGTCGTGGCCTT | RT-qPCR |
| Hsc70-3_R | TTCTTGTACACACCAACGCAGGAAT | RT-qPCR |
| Gp93_F | TTCGCCGGACGAACCGATAGC | RT-qPCR |
| Gp93_R | GGCGGCTATTTGATTGATGCCTGCT | RT-qPCR |
| Xbp1_F | AAGCGTCGCCTGGACCATCT | RT-qPCR |
| Xbp1_R | TCCGTTCTGTCTGTCAGCTCCT | RT-qPCR |
| 7SK_F | GATAACCCGTCGTCATCCAG | RT-qPCR |
| 7SK_R | AGTAATTCTGCCTGGCGTTG | RT-qPCR |
| Xbp1-s-F | CCGAATTCAAGCAGCAACAGCA | RT-PCR for splicing |
| Xbp1-s-R | TAGTCTAGACAGAGGGCCACAATTTCCAG | RT-PCR for splicing |
| Dom-A-RNAi-#2_F | taatacgactcactatagggAGACCAATCAGCCACAACAACAG | dsRNA synthesis |
| Dom-A-RNAi-#2_R | taatacgactcactatagggAGACTGCCTGCACAGTAGTGGAA | dsRNA synthesis |
| Xbp1-RNAi_F | ttaatacgactcactatagggagaCAGCAGCACAACACCAGA | dsRNA synthesis |
| Xbp1-RNAi_R | ttaatacgactcactatagggagaTGTGGGTTTCCATTTATCTTCA | dsRNA synthesis |
| Ctrl(Gst)-F | taaatacgactcactatagggAGAATGTCCCCTATACTAGGTTA | dsRNA synthesis |
| Ctrl(Gst)-Rep | taaatacgactcactatagggAGAACGCATCCAGGCACATTG | dsRNA synthesis |
